# End-to-end bimodal anti-counterfeiting by informational DNA nanoparticles

**DOI:** 10.64898/2026.04.06.716834

**Authors:** Ting He, Bingzhao Zhuo, Xing Zhao, Fukuan Hou, Zhuoming Zhao, Yifei Gong, Jingkang Cao, Lu Liu, Kejing Feng, Ju Tang, Shanni Li, Zhekun Xie, Anni Li, Hui Wang, Qingyang Zhao, Zhunyi Yang, Zhoujun Luo, Zhouqing Luo

**Author notes:** Correspondence (Zhouqing Luo). Co-first author (Ting He, Bingzhao Zhuo, Xing Zhao, Fukuan Hou).

## Abstract

The superior stealth properties and high information density make DNA a sought-after candidate in the field of molecular steganography. Here, we developed the **InfinMark** end-to-end DNA steganography framework for anti-counterfeiting applications by combining the characteristics of both the Internet of Things (IoT) and DNA-of-Things (DoT). InfinMark includes five modules: **In**formation Transcoding, **Fi**ngerprint Writing, **N**ano-encapsulation, Invisible **Mark**ing, and Multi-level Rapid Authentication. It ensures precise anti-counterfeiting information reading and writing through a dynamic DNA-compatible transcoding algorithm, achieves seamless embedding by developing scalable nanoparticle manufacture methods, and supports cross-scenario on-site verification, ultimately granting it comprehensive anti-counterfeiting capabilities spanning from source labeling to terminal tracing. By addressing the bottlenecks in IoT and DoT integration, lifecycle tracking, as well on-site product authentication, this research constructs a full-chain bimodal anti-counterfeiting system, thereby showcasing the practical application of informational DNA nanoparticles in various aspects of production and daily life.

---

The Internet of Things (IoT) has reshaped the operational paradigm of the contemporary world and is poised to play an even more pivotal role in the future realms of artificial intelligence (AI) and automation ^1^. Information carriers such as Universal Product Code (UPC) barcodes and Radio Frequency Identification (RFID) tags, along with their enhanced technologies, act as the intermediaries for information exchange among objects and are integral to the IoT infrastructure. Nevertheless, these carriers inherently suffer from vulnerabilities, including susceptibility to counterfeiting, irreparability following accidental damage, and limited adaptability to substances with unique physical forms. With the progression of DNA information storage technology, a DNA-of-things (DoT) storage architecture has been recently introduced for creating materials with embedded memory ^5^. The potential integration of IoT with DoT to develop informational nanoparticles capable of being incorporated into any object, thereby providing each object with an unforgeable and damage-resistant IoT information carrier credential, will exhibits substantial promise in application areas such as anti-counterfeiting and traceability systems.

Counterfeit products and sophisticated forgery techniques have disrupted markets, causing financial losses for companies and risks to consumers ^1^. While tangible anti-counterfeiting methods like Universal Product Code (UPC) barcodes, Radio Frequency Identification (RFID), and Quick Response (QR) codes, have been standard in commercial anti-counterfeiting and traceability for decades, DNA molecular steganography offers a next-gen alternative with its high information density, invisible nature, and ease of encryption ^2^. However, despite the availability of commercial DNA-tagging services ^3^, adoption of DNA molecular steganography in IoT remains absent due to several unresolved challenges:

1. A major challenge lies in integrating DNA molecular steganography with current standards. On one side, the higher cost and lower throughput of DNA reading/writing technologies hinder real-time tracking of goods circulation data, limiting their application in high-frequency trade, despite the continuous emergence of new approaches ^4^. On the other side, current tangible anti-counterfeiting technologies, predominantly based on surface labels, lack effective solutions for goods with special forms (e.g., fine powders, flexible liquids) or dynamically changing physical states ^2,3^. Mainstream providers currently use DNA taggants as simple identity markers ^3^ with minimal integration with tangible anti-counterfeiting technologies. Thus, a informational DNA nanoparticle based technique combining the Internet of Things (IoT) ^1^ and DNA of Things (DoT) ^5^—while requiring minimal logistics changes—is highly desirable.
2. DNA instability caused by environmental factors (e.g., temperature, humidity, UV light, molecular oxygen, and nucleases) ^2^ necessitates encapsulation in silica ^6^ microbial spores ^7^ or polymers ^8^ to create protective barriers against degradation for lifecycle tracking. However, the use of hazardous reagents like fluoride-containing or organic solutions to prepare or dissolve these barriers makes large-scale manufacturing impractical, restricting applications to controlled laboratories. It is necessary to develop non-toxic and cost-effective large-scale DNA nanoparticle preparation methods.
3. On-site authentication remains a bottleneck, as current methods rely on specialized PCR protocols and laboratory equipment ^9^, hindering field deployment. The key to widespread adoption lies in developing cost-effective, lab-free on-site authentication methods.

## InfinMark: an integrated molecular steganography strategy

To address the aforementioned challenges and equip DNA with molecular steganography capability that spans from source labeling to terminal tracing, we established InfinMark - an end-to-end anti-counterfeiting and traceability framework that unifies five core processes (Figure 1): **In**formation transcoding, **fi**ngerprint writing, **n**ano-encapsulation, invisible **mark**ing, and multi-level rapid authentication.

**Figure 1.**
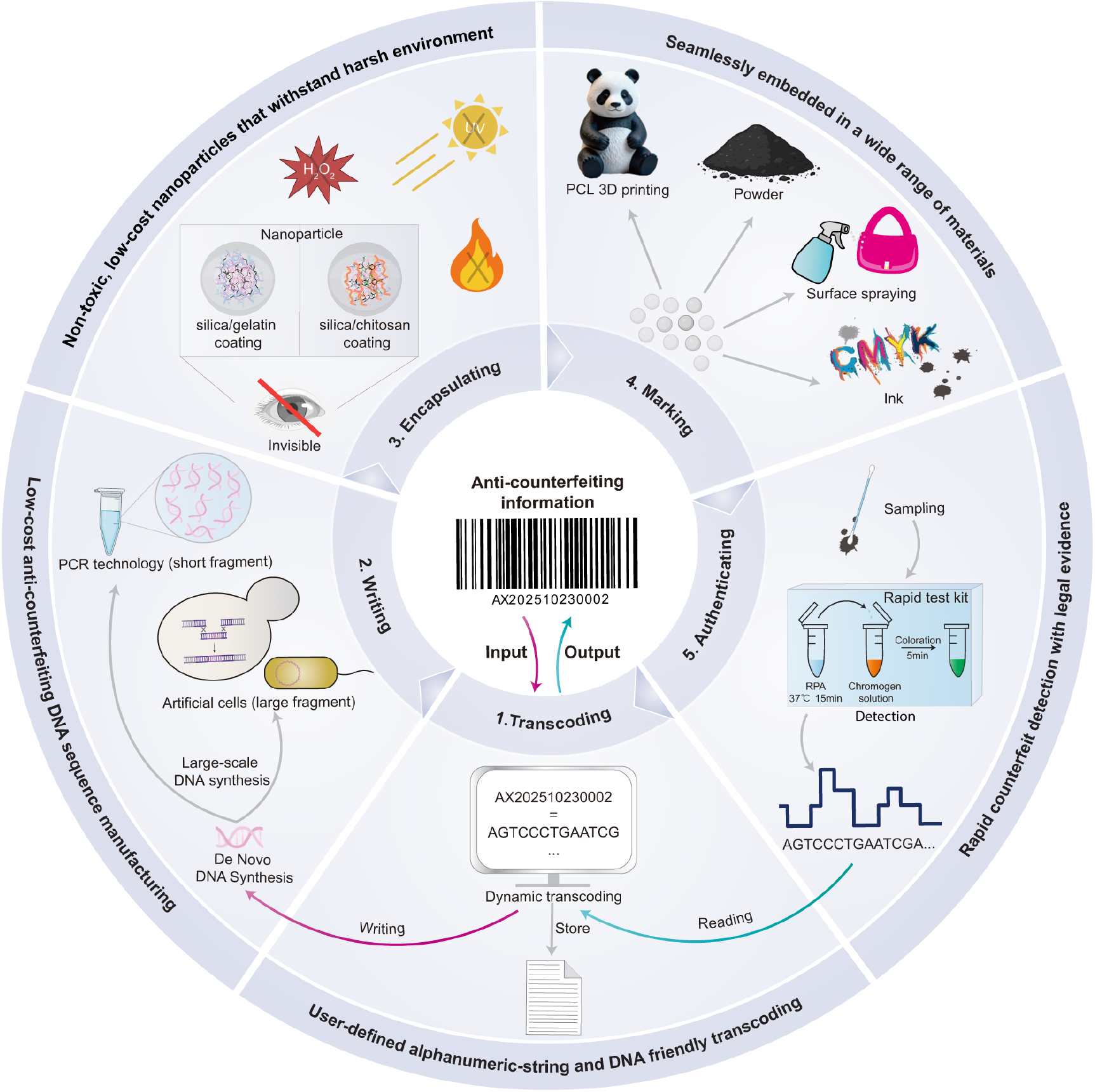
Creating a holistic product anti-counterfeiting and traceability framework across the entire chain utilizing DNA-based cryptographic conversion. A novel dynamic transcoding algorithm enables bidirectional, encrypted conversion between anti-counterfeiting information and DNA sequences, facilitating ‘one-click’ encoding and decoding for users. The DNA sequences are then converted into DNA fingerprints via *de novo* synthesis, followed by scalable amplification through PCR or microorganisms depending on the required complexity. These DNA fingerprints are encapsulated through different methods within a dual-layer protective barrier, which ensures long-term stability and integrity from production to end-use by withstanding environmental stressors such as UV radiation, H_2_O_2_ exposure, and freeze-thaw cycles, etc. The resulting nano-level, invisible DNA nanoparticles can be embedded into various materials forms (powders, liquids and solids) without altering their properties, enabling anti-counterfeiting at the source level. An InfoTrace kit was developed for on-site authentication in multiple scenarios and provides forensic-grade evidence when used in conjunction with sequencing.

1. **Information transcoding**. To ensure integration with tangible anti-counterfeiting technologies, we developed a **dy**namic **t**ranscoding **a**lgorithm called DYTA. DYTA enables the “one-click” conversion of alphanumeric-string-based information—which is widely utilized in mainstream technologies like barcodes, QR codes, and blockchain—into DNA sequences, while ensuring cryptographic robustness. This has established a dual-coding (overt and covert) anti-counterfeiting system, where tangible systems like UPC are employed for real-time tracking of goods circulation data, and DNA serves as the core cryptographic key for authenticating these information.
2. **Fingerprint writing**. By rigorously optimizing key parameters that govern DNA synthesis and detection efficiency, such as GC content and homopolymer length, DYTA produces virtual DNA sequences that are both highly amenable to efficient synthesis and significantly enhance detection accuracy and throughput. The extremely high information density effectively reduces the length of DNA that needs to be synthesized. DNA sequences are first converted into DNA fingerprints in small quantities via *de novo* DNA synthesis, then economically amplified by PCR and assembled by *E. coli* or yeast.
3. **Nano-encapsulation**. A nano-encapsulation strategy was implemented that isolates the DNA within a dual-layer protective barrier, thereby ensuring its lifecycle stability and integrity. The use of food-grade preparation materials and a simplified preparation process enables large-scale production and simple extraction.
4. **Invisible marking**. Inspired by the DoT concept ^5^, the diameter of the encapsulated DNA nanoparticles were precisely controlled at the nanoscale, and the nanoparticles were then embed in trace amounts into various materials, including inks, powders, and solids, without altering their properties. This approach provides a source-level anti-counterfeiting solution that is highly resistant to forgery for products incompatible with conventional label-based methods.
5. **Multi-level authentication**. A field-deployable InfoTrace kit was introduced to achieve pg-level molecule detection sensitivity. The entire procedure requires only minutes and a simple protocol, analogous to a standard COVID-19 test, thereby eliminating the dependency on specialized equipment and expertise. Following initial detection, the choice of sequencing method can be tailored to the scale of the anti-counterfeiting DNA samples and the user’s urgency for results, thus providing precise, forensics-level evidence for anti-counterfeiting and traceability.

## Dynamic transcoding algorithm compatible with alphanumerics and DNA

An end-to-end isolated architecture is employed for information transcoding and transmission (Figure 2A). During encoding, information is transcoded into DNA fingerprints at the local client, with the encoding rules stored in a locally encrypted codebook to prevent interception. Subsequently, these DNA fingerprints are transferred to an internet-connected computer via an offline medium such as a USB drive and then sent to DNA synthesis providers. During decoding, after receiving the sequenced DNA data, this data is similarly transported back to the local client through a secure medium and decoded into the original anti-counterfeiting information based on the pre-stored codebook. Throughout the entire transcoding workflow, all external entities, including DNA synthesis providers, sequencing facilities, and even anti-counterfeiting product manufacturers, are limited to accessing only the encrypted DNA sequences. This architecture inherently blocks unauthorized access to core cryptographic modules and keys, ensuring procedural security and maintaining strict information neutrality across the entire information processing chain.

**Figure 2.**
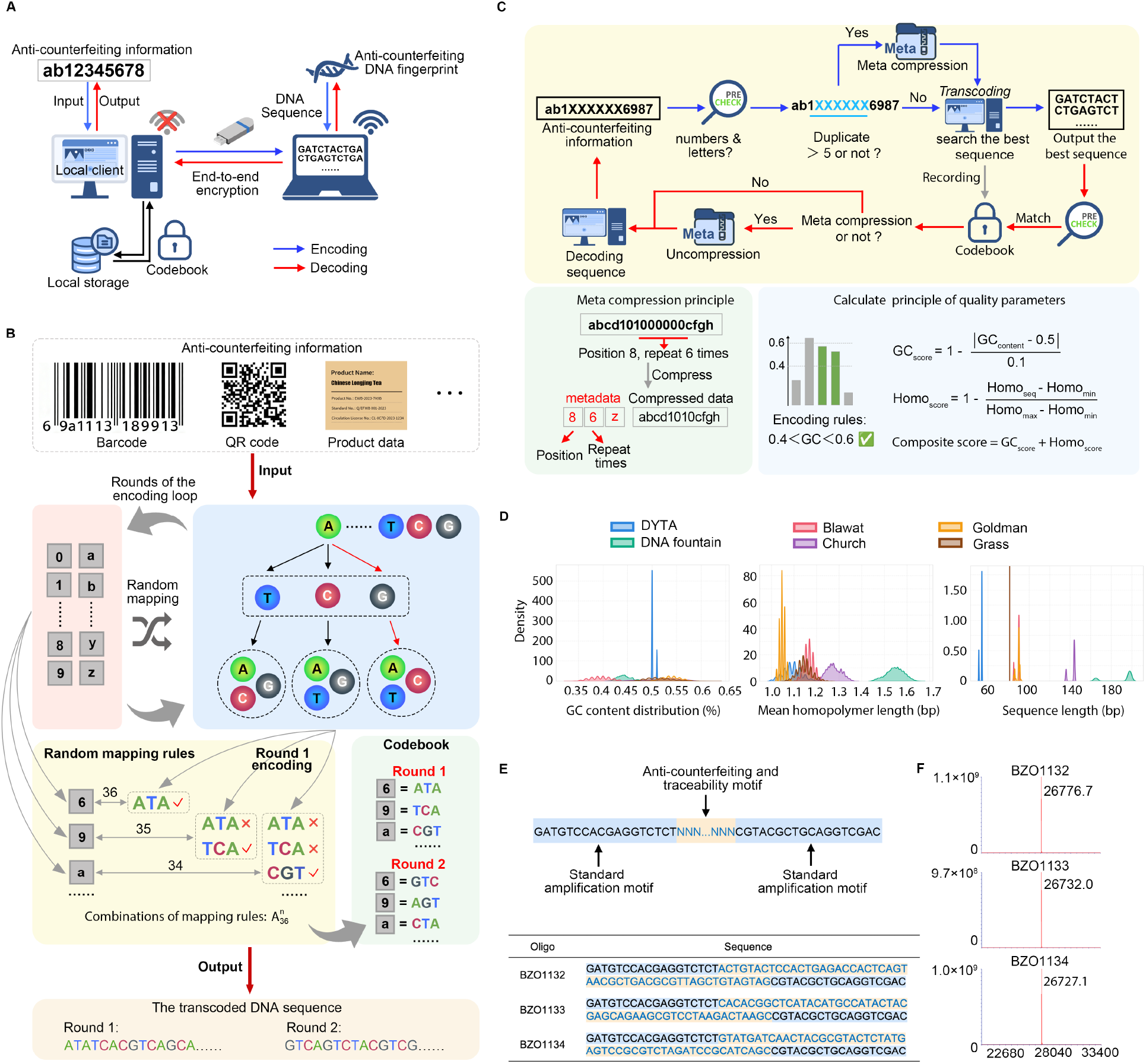
Dynamic Encoding and Decoding Module. **A**. Local encoding and decoding system. Anti-counterfeiting and traceability data is encoded using an offline host, transferred via portable storage to an online terminal for synthesis, and stored locally along with the codebooks. For decoding, sequencing DNA data is returned through the same secure path to reconstruct the original authentication information. **B**. Mechanism of DYTA dynamic encoding and decoding algorithm. **C**. Workflow for processing authentication information containing more than five consecutive repeated characters, including meta-compression, encoding/decoding, and the application of scoring and selection criteria to output sequences. **D**. Comparison of DNA sequences generated by Blawat, Goldman, DNA Fountain, Church, Grass, and DYTA. **E**. Composition of anti-counterfeiting DNA sequences. The structure consists of a central traceability information segment, generated through transcoding, which is flanked by universal standard amplification sequences at both ends. **F**. The three DNA sequences with the lowest scores from the dynamic encoding pool were selected for synthesis and validated by mass spectrometry.

The core of this architecture lies in the dynamic codec called DYTA designed for seamless transcoding between anti-counterfeiting information and DNA sequences. For alphanumeric-string-based information, which is widely applied in mainstream technologies such as barcodes, QR codes, and blockchain, DYTA uniquely maps each of the 36 alphanumeric characters to one of 36 different three-base combinations (with 4×3×3 possible combinations, excluding adjacent identical bases). This scheme theoretically enables a code space up to 36 factorial (36!). Taking the barcode “69a1113189913” shown in Figure 2B as an example, the encoding process proceeds sequentially: the first character ‘6’ can be mapped to any of the 36 three-base combinations, the second character ‘9’ can be mapped to any of the remaining 35 combinations, and so forth. In general, for information containing n distinct characters, the total number of possible unique mapping rules is the permutation number 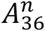. This design can prevent the occurrence of three consecutive identical bases in the generated DNA sequence, thereby reducing the formation of secondary structures, enhancing resistance to synthesis and sequencing errors, and facilitating subsequent molecular manipulations (such as assembling large DNA fragments encoding a vast amount of information).

Importantly, each encoding round employs a stochastic and independent set of keys. For example, the same plaintext “69a11…” may be translated into sequence “ATATCACGTCAGCA…” in one round, and into “GTCAGTCTACGTCG…” in another. This not only mitigates pattern-based attacks but also endows the scheme with the cryptographic strength of a one-time pad. This dynamic encoding algorithm ensures identical input data produce distinct nucleotide sequences across different sessions, thus mapping various anti-counterfeiting identifiers (such as barcodes and QR codes) and product metadata (including blockchain transaction hashes) into customized DNA sequences.

Consecutive identical characters in the source data form repetitive DNA sequences of multiple repeated base triplets upon encoding. To mitigate this, we introduced a metadata-based compression scheme for anti-counterfeiting information with exceeding 5 identical characters (Fig. 2C, Fig. S1A), wherein the repeat position (addr) and length (data) are extracted to form a metadata sequence (meta_seq) for subsequent transcoding. For instance, the string “abcd101000000cfgh”, containing six consecutive ‘0’s starting at position 8, is compressed into “86z”“abcd1010cfgh” (Fig. 2C).

To fine-tune the DNA sequence within an optimal GC content range (0.4–0.6) and minimize homopolymer formation, thereby improving synthesizability, we implemented a dynamic mapping strategy by rotating the assignment between characters and DNA coding sets. Candidate mapping rules that met the GC content criteria were further evaluated using a dual-parameter scoring system incorporating both GC score and homopolymer score (Fig 2C). The sequence with the highest total score was prioritized for synthesis; in cases of tied scores, preference was given first to the sequence with a higher GC score, followed by a higher homopolymer score (Fig. 2C, Fig. S1A). For the decoding workflow (Fig. 2C, red arrow; Fig. S1B), the input DNA sequence is validated and matched to a codebook. Whenever a metadata identifier is detected, decompression is performed using the metadata to restore the complete original DNA sequence.

To validate performance at scale, we converted 500 MB of real-world tea anti-counterfeiting data into DNA sequences (see Methods). Our experimental results demonstrate the exceptional suitability of DYTA in anti-counterfeiting applications (Fig. 2D, comprising a total of 19,951 barcodes). The generated DNA fingerprints exhibit near-ideal GC content (50%), a low average homopolymer length (1.1 base pairs), and a minimum encoded sequence length of only 51 base pairs. This indicates that such sequences can be synthesized, sequenced, and mass-produced at low cost through commercial DNA synthesis and sequencing platforms and are compatible with industrial manufacturing processes. Mass spectrometry analysis of the three lowest-scoring DNA sequences confirmed that their molecular mass deviations were all ≤ 0.05%, highlighting the exceptional performance of DYTA in precisely controlling the characteristics of DNA sequences (Fig. 2E-F, Table S1).

## Informational DNA nanoparticles for lifecycle authentication

DNA fingerprints’ susceptibility to harsh conditions (UV, ROS, extreme temperatures, etc.) poses a significant challenge, limiting their practical use in anti-counterfeiting. To safeguard DNA throughout a product’s lifecycle, we propose a hierarchical dual-encapsulation strategy that combines a molecular network with a mineral shell (Fig. 3A). For the first defense layer, we construct a 3D network by crosslinking chitosan with vanillin, in parallel with employing the gelatin-glutaraldehyde strategy ^10^. To further bolster stability and compatibility, sodium silicate is deposited onto the DNA-loaded 3D network via charge-mediated interactions. This process results in the creation of impermeable silica nanoparticles, offering complete isolation and safeguarding DNA fingerprints from external environmental disturbances and degradation.

**Figure 3.**
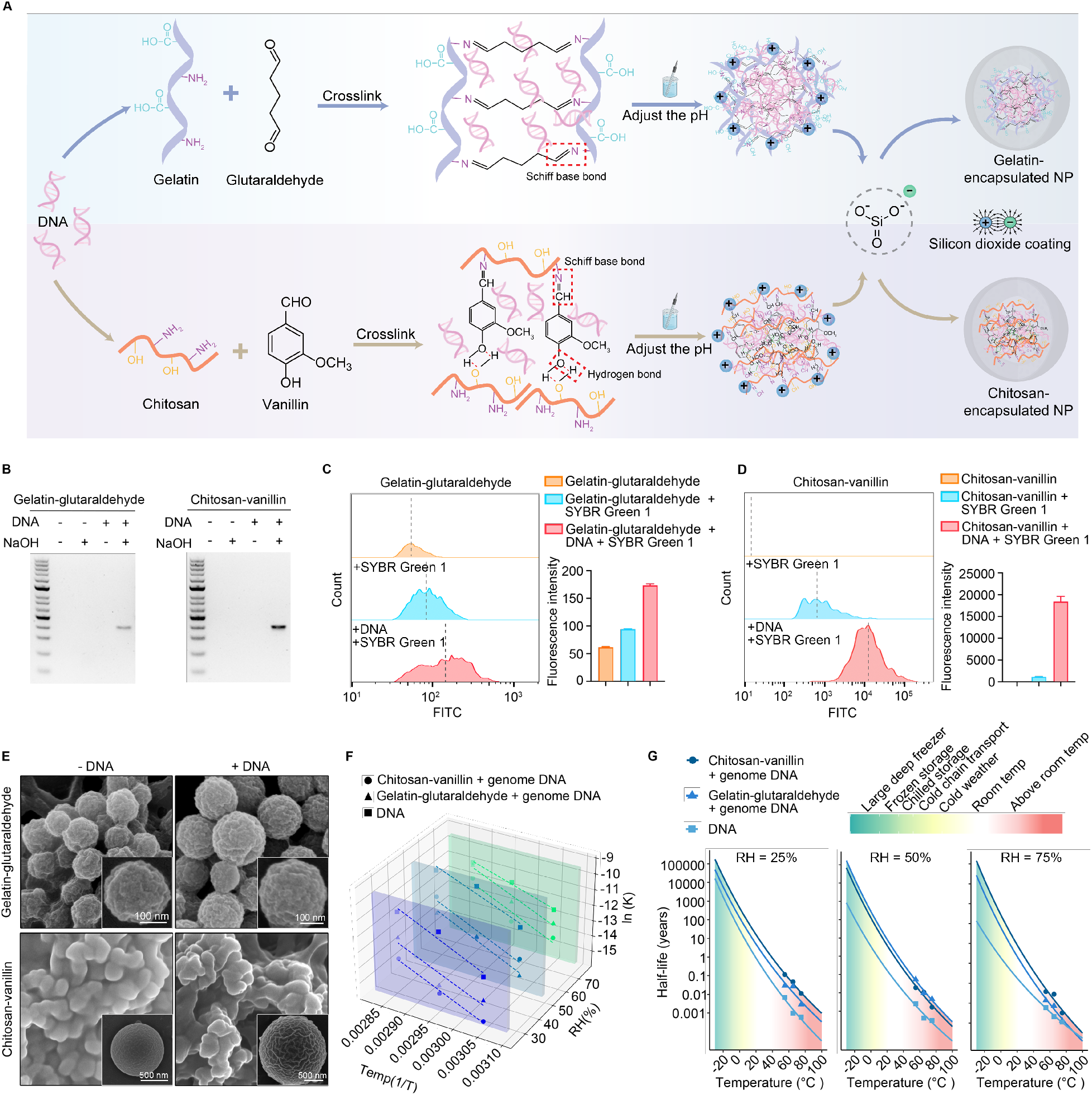
Preparation and performance of infomatic DNA nanoparticles. **A**. Principle and flowchart of DNA fingerprint encapsulation. Under acidic conditions (pH 3.2), the negatively charged DNA first interacts with positively charged gelatin or chitosan via electrostatic attraction, and the complexes are then cross-linked with glutaraldehyde or vanillin, respectively. The polymerization degree is controlled by pH adjustment, after which sodium silicate is added to form a silica shell around the complexes. **B**. Analysis of DNA fingerprints post-encapsulation. Following DNase I digestion, encapsulated DNA fingerprint nanoparticles, with or without NaOH decapsulation, were analyzed by PCR and gel electrophoresis. **C-D**. Verification of DNA fingerprint encapsulation by Nano-flow Cytometry. The following negative controls were established: (1) gelatin-glutaraldehyde or chitosan-vanillin encapsulation particles without DNA, and (2) particles encapsulating SYBR Green 1 dye alone. The bar chart represents the mean fluorescence intensity of each group, presented as mean ± S.E.M. **E**. SEM images of DNA fingerprint NPs and the morphology of representative individual particles. Scale bar for gelatin-glutaraldehyde: 100 nm, and Scale bar for chitosan-vanillin, 500 nm. The aggregation observed in the chitosan-vanillin group under SEM is likely an artifact of sample preparation, as no aggregation was detected in solution or during subsequent applications. **F**. Degradation kinetics of dried DNA fingerprint NPs under varying temperature and humidity. Arrhenius plots (ln(k) vs. 1/T) for both naked and encapsulated DNA in either gelatin-glutaraldehyde or chitosan-vanillin matrices exhibited mutually parallel linear relationships across the three humidity conditions. **G**. The half-lives of naked and encapsulated DNA fingerprints were calculated according to the Arrhenius equation. RH, relative humidity.

To evaluate the encapsulation efficacy, all the samples were treated with DNase I to digest any unencapsulated DNA, followed by extraction of the encapsulated DNA using mild 0.2 M NaOH with heat cycling (see Methods), a gentler alternative to conventional harsh hydrogen fluoride or concentrated alkali solution ^6^. The subsequent positive PCR results in the NaOH treatment group confirm both an effective encapsulation/extraction process and robust protection against enzymatic degradation (Fig. 3B). Pre-encapsulation labeling of DNA with SYBR Green I enabled direct comparison of DNA loading capacity via nano-flow cytometry (Fig. 3C–3D). The chitosan-vanillin nanoparticles exhibited a 100-fold higher fluorescence intensity than the gelatin-glutaraldehyde group under identical conditions, indicating superior DNA loading. SEM and size distribution analyses also revealed distinct structural differences between the two encapsulation systems. The gelatin-glutaraldehyde system self-assembled into monodisperse spherical nanoparticles with a highly narrow size distribution [average diameter: 338 nm; polydispersity index (PDI): 0.076], indicating higher homogeneity (Fig. 3E, Fig. S2A). Meanwhile, the chitosan-vanillin system formed moderately polydisperse colloidal particles (mean diameter: 420.4 ± 12.59 nm; PDI: 0.224), and defined spherical morphologies were observed at low concentrations (Fig. 3E, Fig. S2B). Zeta potential measurements revealed distinct surface charge profiles for the two systems, with gelatin-glutaraldehyde NPs possessing a negative charge and chitosan-vanillin NPs a positive charge (Fig. S2C-D). The distinct characteristics of the two materials suggest their potential division across different application niches and specific electrical environments.

To systematically evaluate how encapsulation and extraction affect DNA sequence fidelity, we employed anti-counterfeiting DNA nanoparticles containing the Xiamen University motto “zi qiang bu xi zhi yu zhi shan” as a model. Sequencing results demonstrated a minimum single-base accuracy of 98.6% across all conditions without correction of intrinsic sequencing errors and showed no statistically significant discrepancy from direct sequencing of raw DNA fingerprint amplicons (Fig. S3A). Notably, original sequences could be reliably recovered at mean 2× sequencing depth, while sequences harboring mutations at the same position required only 5× depth for complete reconstruction (Fig. S3B, Table S2). At a 5× sequencing depth, the probability that the reconstructed sequence based on sequencing results differs from the original one will be less than 0.0001 (FDR = 0.01389) (Table S2). From an application perspective, this low depth requirement enables high multiplexing (i.e., more samples per sequencing run) on platforms like Illumina, dramatically lowering the per-sample sequencing cost and improving overall cost-effectiveness.

To systematically assess the stability and recoverability of encapsulated DNA fingerprints under harsh conditions, including UV irradiation, freeze-thaw cycles, and hydrogen peroxide exposure, a series of tolerance tests were conducted (see Methods). As shown in Fig. S4A, the gelatin– glutaraldehyde group retained 50% of DNA after 2 hours of UV irradiation, and the chitosan– vanillin group maintained over 50% throughout the 4-hour test, whereas naked DNA dropped to only 6% within 1 hour. This significantly enhances protection against UV-induced DNA photodegradation, a key weakness of conventional silica encapsulation due to the material’s inherent transparency ^6^. In freeze-thaw tests (Fig. S4B), encapsulated nanoparticles preserved DNA integrity completely over 30 cycles, compared to merely 1.5% retention for naked DNA. Under oxidative stress, naked DNA retention was reduced to 2% following 5 mM H_2_O_2_ treatment, while the gelatin-glutaraldehyde encapsulated DNA retained nearly 8% even after >30 mM H_2_O_2_ treatment, and the chitosan-vanillin group maintained over 18.7% under the same conditions (Fig. S4C). Both encapsulation methods significantly reduced the overall mutation frequency upon UV irradiation (Fig. S5A, S5D), and the chitosan-vanillin group also exhibited a protective effect under freeze-thaw cycling and hydrogen peroxide treatment (Fig. S5B–D). It should be noted that the sequencing library preparation relied on successful PCR amplification of post-treatment samples. Consequently, mutations that prevented amplification, such as key point mutations in the primer-binding region, large insertions/deletions, or the formation of complex secondary structures, could not be assessed. This limitation may lead to an underestimation of the true differences in protective efficacy among the encapsulation groups.

Even under the above aggressive conditions, the original anti-counterfeiting information remains fully recoverable. Importantly, the DNA sequences of encapsulation groups could be accurately recovered using only 3× sequencing depth, significantly outperforming the naked DNA particularly when exposed to UV conditions (Fig. S6). These data suggest that the nanoparticles provide both elasticity and mechanical strength to buffer external physical impact and fluid shear stress, thereby preventing mechanical breakage of DNA strands. The dense structure of the nanoparticles serves as a physical barrier against external harmful agents, and their colloidal nature helps maintain a stable internal microenvironment, slowing chemical hydrolysis.

To evaluate the long-term stability of the anti-counterfeiting DNA nanoparticles under normal storage conditions, we referred to established accelerated aging experimental methods ^11^. The results showed that under different humidity conditions, both the unencapsulated DNA powder and the encapsulated nanoparticles exhibited a linear relationship between ln(k) and 1/T (where k is the first-order decay rate constant and T is the temperature in Kelvin), with fitted lines that were parallel, indicating essentially the same activation energy for the degradation reaction (Fig. 3F). This suggests that encapsulation leaves the chemical energy barrier for DNA degradation unchanged but operates at the molecular dynamics level by limiting the accessibility and diffusion between reactants, thereby decreasing the effective collision frequency, lowering the DNA decay rate (k), and ultimately protecting the DNA.

Based on the Arrhenius equation ^11,12^, the half-life of the DNA fingerprints in the two encapsulation methods significantly increased with decreasing temperature and humidity, consistently surpassing that of unencapsulated DNA (Fig. 3G). Extrapolation of the data to common storage and transportation conditions predicts greatly extended stability (Table S3). Under typical storage and transportation conditions, the projected preservation times far exceed practical needs: several decades at room temperature and centuries under refrigeration. It is important to note that predictions for extreme conditions (e.g., cryogenic storage) represent theoretical extrapolations based on the model and should be interpreted with caution, as long-term stability at such temperatures would require empirical validation.

## On-site Multi-Scenario Authentication

Seamless embedding and rapid detection are two core requirements for the practical application of DNA anti-counterfeiting. The nanostructure endows the DNA fingerprint with stability and material compatibility, protecting the DNA not only from material-induced degradation but also from damage caused by morphological changes and physical stresses during product processing. Owing to its nanoscale size and invisible nature, this technology overcomes the size limitations inherent to conventional labels. Unlike surface-applied tags, it allows covert embedding of anti-counterfeiting tags directly into the material matrix, whether solid, liquid, or powder, effectively preventing tampering or removal, establishing an invisible anti-counterfeiting technique (Fig. 4A). To enable rapid on-site product authentication, a pg-level detection process based on Recombinase Polymerase Amplification (RPA) was developed (see Methods and Fig. S7). Simple alkaline lysis followed by 5–15 minutes of isothermal amplification is enough to generate the amplicon, which is then diluted and mixed with a detection reagent to yield a visible color change for authentication (Fig. 4A, Fig. S8). Furthermore, when combined with high-accuracy sequencing, the results can be certifiable for legal verification and provide court-acceptable documentation of product provenance (Fig. 4A and 4B). This instrument-free method is suitable for on-site authentication applications such as origin verification of products from incident scenes or supply chain spot checks.

**Figure 4.**
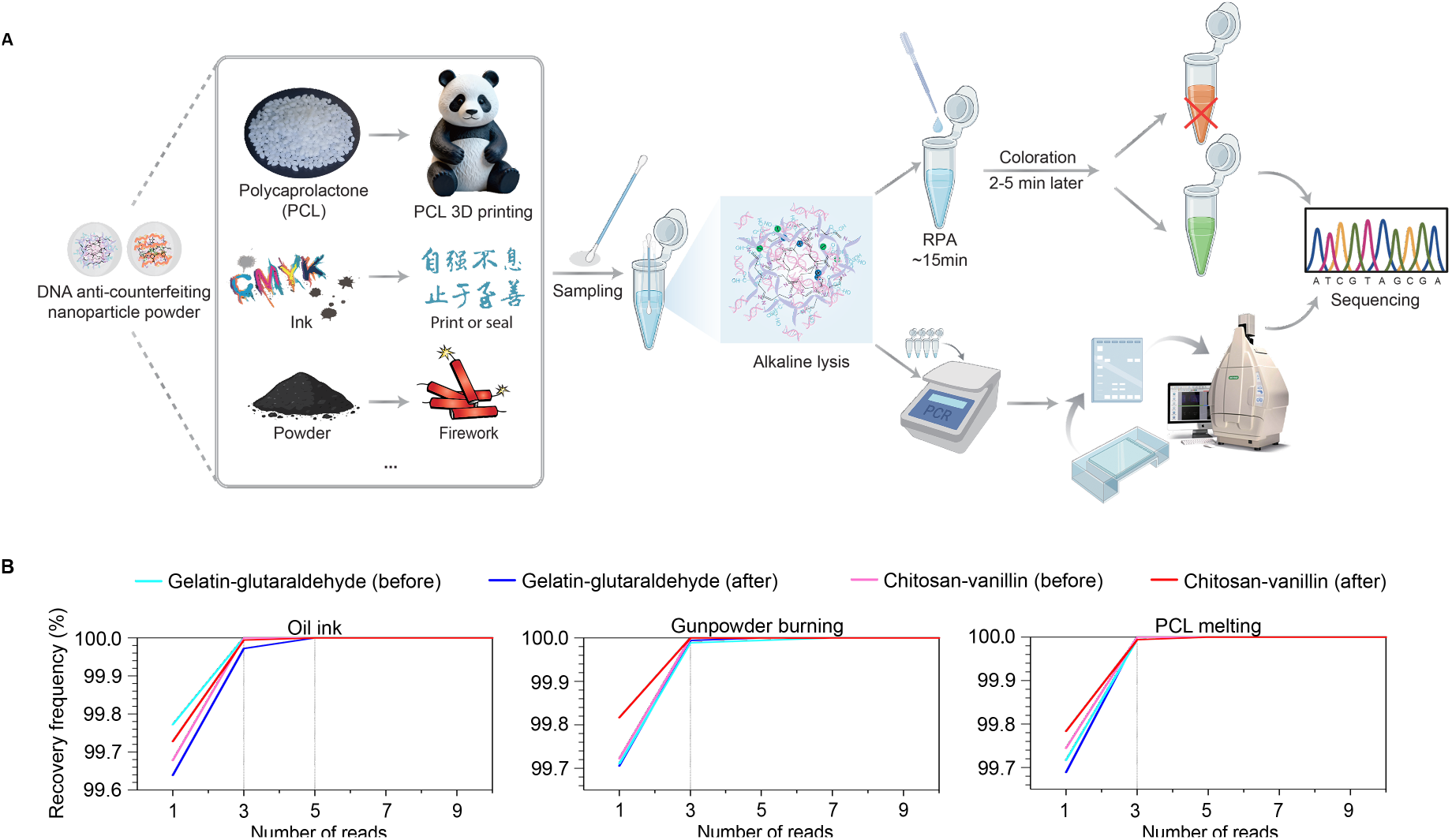
Application and readout of covert anti-counterfeiting DNA fingerprint NPs. **A**. Schematic diagram of anti-counterfeiting and tracing using DNA fingerprint NPs in plastics, oil inks, and gunpowder. **B**. DNA fingerprint NPs were extracted from stamped ink on paper, combusted gunpowder residue, and melted polycaprolactone (PCL), followed by amplification and sequencing. The minimum sequencing depth required for accurate recovery was determined based on the Illumina sequencing data.

To evaluate the practical stability and compatibility of anti-counterfeiting DNA fingerprint nanoparticles, we incorporated them at a 1:500 (w/w) ratio into three representative material matrices: ink (liquid), gunpowder (powder), and bioplastic PCL (polycaprolactone) (Fig. S8). Rather than replacing conventional marking technologies such as QR codes, the embedding of DNA nanoparticles into ink is designed to establish a “dual-code” authentication system. While the visible QR code maintains its functions in rapid reading, blockchain registration, and circulation tracking, meeting routine verification and digital management needs, the hidden DNA fingerprint acts as an imperceptible authentication anchor embedded within the material itself. Even under conditions of physical wear, partial contamination, or intentional damage, the DNA nanotags remain extractable and verifiable from residual ink traces, effectively overcoming the inherent vulnerabilities of surface labels to peeling, transfer, or tampering. In a practical demonstration, a seal bearing the Xiamen University motto “ 自强不息, 止于至善 ” was used to stamp paper with InfinMark ink (Fig. S8, left; see Supplementary Video, 0:10 **–** 0:50). After the imprint had fully dried, samples were collected from both the original oil ink and the dried paper imprint. The target DNA signal was consistently detected in both samples using PCR and our InfoTrace kit (Fig. S8, left; see Supplementary Video, 2:21 **–** 4:26). Furthermore, the close match in key physical parameters, such as particle size and PDI, between the anti-counterfeiting nanoparticles and conventional printing inks ensures seamless integration into modern printing systems.

The authentication and traceability of powdered hazardous materials such as gunpowder remain a critical challenge, constrained both by the size limitations of conventional labeling methods and and by stringent demands for quality control and illicit trafficking suppression. To address this, we incorporated DNA nanoparticles into gunpowder and evaluated their stability under deflagration conditions (see Supplementary Video, 0:51 **–** 1:35). The resulting InfinMark gunpowder exhibited undiminished ignition capability (Fig. S8, middle). Subsequent analysis using both an InfoTrace kit and PCR confirmed the presence of DNA fingerprints in post-deflagration residues, as evidenced by a distinct green color from the chromogenic reaction and a clear positive PCR band (Fig. S8; see Supplementary Video, 2:21 **–** 4:26). These signals were fully consistent with those from pre-combustion samples, confirming the high compatibility and stability of the DNA anti-counterfeiting nanoparticles within the powder matrix.

Meanwhile, authenticating objects that undergo physical changes presents another formidable challenge for conventional tagging systems. The biodegradable PCL plastic, for example, is a polymeric solid material that undergoes such physical transformations and is widely used in packaging, medical devices, and 3D printing. In practical applications, PCL often undergoes melting and solidification cycles, requiring anti-counterfeiting DNA NPs to maintain stability within a dynamically changing matrix-a demanding task for reliable authentication. In this study, DNA NPs were blended with PCL particles and subjected to a thermal cycle involving melting at 100°C followed by solidification at room temperature. The InfinMark remained invisible in both the transparent molten and the re-solidified states (Fig. S8, right; see Supplementary Video, 1:36 **–** 2:20). Notably, DNA extracted from the re-solidified samples generated unambiguous green signals when analyzed with the InfoTrace kit (see Supplementary video, 2:21 **–** 4:26), a result consistent with the pre-melting samples and corroborated by PCR amplification (Fig. S8, right). This confirms the robustness of the InfinMark-based authentication system for materials that undergo physical changes throughout their lifecycle in application.

Amplicons derived from samples collected before and after the processes of stamping, gunpowder deflagration, and PCL melting were sequenced on the Illumina platform. Across all samples, the the intended target DNA sequences were consistently recovered at an average sequencing depth of 5×, with a sequence reconstruction error probability below 0.0001 (FDR < 0.001) (Fig. 4B, Fig. S9, Table S2)

## Technical and Economic Analysis

The widespread adoption of DNA-based anti-counterfeiting technologies depends critically on their technical maturity and market competitiveness. To systematically evalute the industrialization prospects of InfinMark, we compared it with existing research across multiple dimensions, including cryptographic performance, production cycle, ease of use, and environmental friendliness (see Table S4).

For encryption, InfinMark features a dual-security architecture that integrates the inherent complexity of DNA materials with a dynamic encryption algorithm. This design provides superior cryptographic strength and resistance to reverse engineering compared to techniques relying on natural DNA or static coding, achieving a security level approximating that of physicochemical unclonable technologies (Fig. 5A, Table S4).

**Figure 5.**
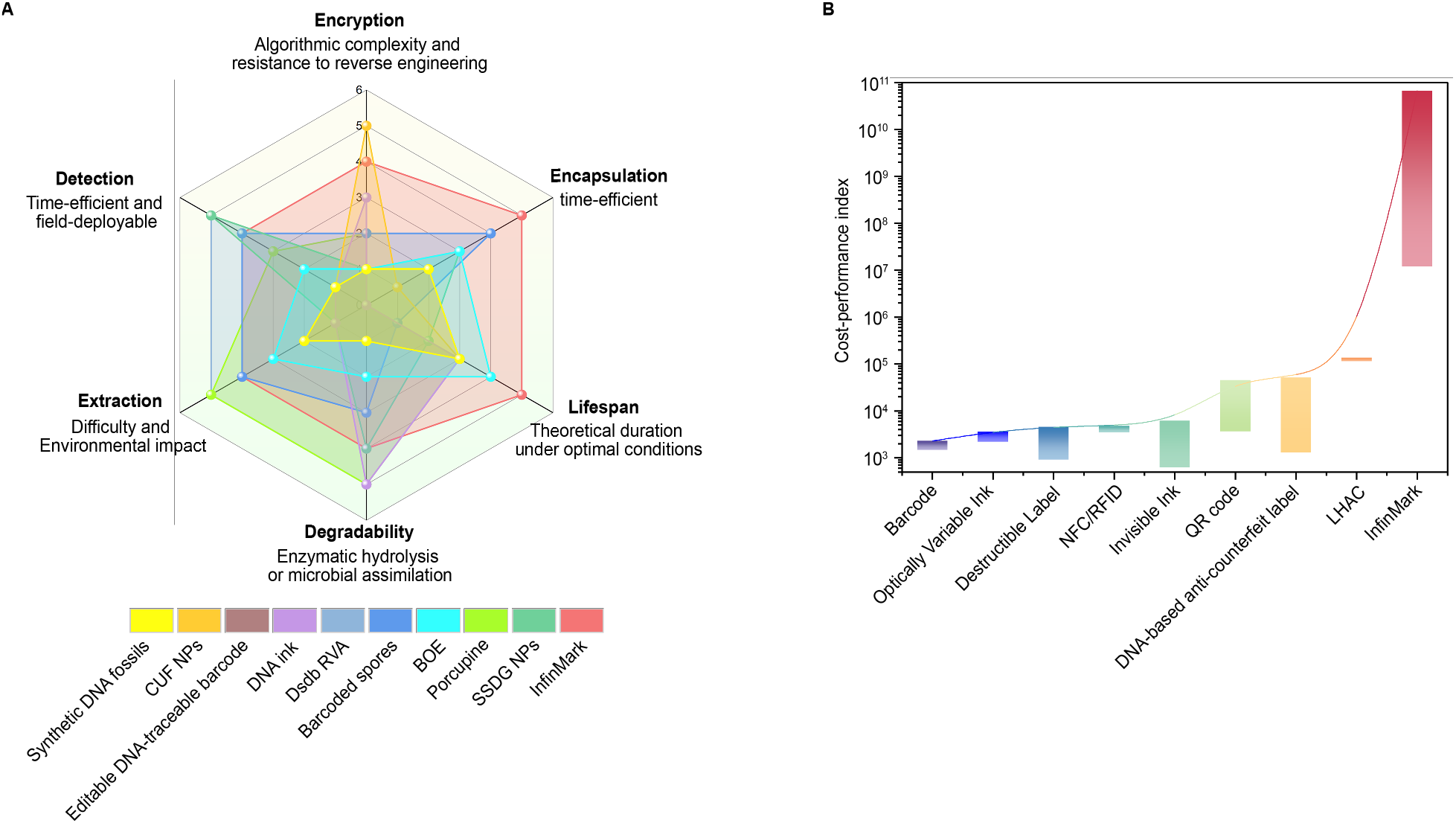
Comprehensive evaluation of Infinmark and mainstream anti-counterfeiting technologies. **A**. Performance comparison based on multiple parameters. The encryption complexity and resistance to reverse engineering, encapsulation duration, nanoparticle lifespan, degradability, complexity of DNA extraction, detection convenience and reagents environmental friendliness were used as comparative parameters. A higher value corresponds to better performance for each metric. Detailed parameter definitions and grading criteria are provided in Table S4 (worksheet “Comparison & Evaluation”). **B**. Cost-effectiveness comparison. Mainstream and emerging methods were compared using a cost-effectiveness index (V), which integrates four parameters: encryption space size, particle lifespan, engineering-experience-weighted encryption level, and unit cost. Detailed data and calculations are in Table S6. The comparative parameters were identified based on data from previous studies and reasonable inference drawn from the practical implementation performance of relevant technologies. SSDG NPs, silica-coated SYBR Green I (SG)/DNA gelatin nanoparticles; Dsdb RVA, DNA-strand-displacement-based rapid visual authentication; CUF NPs,chemical unclonable functions (CUFs) NPs; BOE, biodegradable organosilica encapsulates; LHAC, laser holographic anti-counterfeiting.

In terms of encapsulation and extraction, the encapsulation process of mainstream inorganic silica-based methods requires at least 6 days and depends on toxic extraction reagents, with poor environmental degradability, impeding its practical adoption and ecological compatibility ^5,6,13^. While subsequent refinements have reduced the production cycle to four days and improved certain aspects of environmental compatibility ^7,14,15^, fundamental systemic constraints persist. By contrast, InfinMark employs a bio-based composite encapsulation system (1.5 days), providing superior stability with a lifespan exceeding that of most DNA-based anti-counterfeiting products at ambient temperature (15°C), far surpassing the typical product protection period of >10 years ^14^ (Fig. 5A, Table S4). Moreover, it employs degradable encapsulation materials and an environmentally compatible extraction process using low-concentration NaOH, easily neutralized into non-toxic salts. This approach offers a distinct advantage over the traditional hydrofluoric acid/phenol-chloroform method, providing a greener pathway for scalable industrial manufacturing and applications in sensitive fields like food and pharmaceuticals (Fig. 5A, Table S4). Beyond this benefits, InfinMark exhibits broad material compatibility, enabling integration with materials spanning diverse physical states such as inks, powders, and polycaprolactone (PCL), supporting expanded applications and device integration such as functional printing, specialized energetic materials, and biomedical scaffolds. Collectively, these characteristics allow InfinMark to achieve plug-and-play integration without modifying existing production lines, while ensuring complete extraction of encrypted DNA signatures. It provides a systematic solution for high-security, long-lifecycle, green, and sustainable anti-counterfeiting needs.

As for authentication, current DNA-based methods typically rely on multi-hour PCR/qPCR protocols, which are impractical for on-site use. While fluorescent indicator co-encapsulation enables second-level readout, it drives up production costs and compromises stability, limiting the shelf life at ambient temperatue to approximately one to two years, as seen in examples like SSDG NPs and DNA-Strand-Displacement-based Rapid Visual Authentication ^15,16^ (Fig. 5A, Table S4). Through a systematic comparison of key metrics, including time for authentication, field deployability, and dependence on specialized instruments and personnel, this study demonstrate that InfinMark achieves minute-level authentication using a field-ready kit without compromising long-term stability or requiring advanced equipment or trained operators (Fig. 5A). These combined features give InfinMark a practical edge in overall feasibility among DNA-based anti-counterfeiting approaches.

Under the prevailing dilemma where mainstream solutions lack sufficient security and durability, while high-end alternatives are hindered by cost and complexity ^3^ (Table S5), the future competitiveness of InfinMark lies in its systematic integration of high security, rapid verification, and long-lasting authentication within a controllable cost structure. A detailed cost analysis of materials, reagents, and sequencing shows that the per-tag cost for InfinMark is approximately 0.017–0.028 US cents (Table S6), with encoding costs negligible. Decoding at 5× sequencing depth adds only about 0.0015 cents per tag. When paired with a rapid detection kit, the estimated cost per reaction is around $0.886 based on current commercial reagent pricing, a cost already lower than that of large-scale-procured COVID-19 rapid tests. These estimates are derived from small-scale experiments; scaling up production and detection is projected to substantially reduce costs, strengthening the economic rationale for commercial adoption. To enable a more detailed and objective evaluation, this study further introduced a cost-effectiveness index (V) that incorporates four key dimensions: encryption-space complexity, durability, engineering-experience-weighted encryption level, and cost (see Methods). Results show that InfinMark achieves a significantly higher V-value than both current mainstream anti-counterfeiting technologies and emerging molecular-based solutions (Fig. 5B, Table S6), establishing a decisive lead in overall cost-effectiveness.

## Discussion

The practical application of synthetic DNA tags in the market remains limited nearly four decades after the concept was first introduced ^3^. The core challenge stems from the significant practical constraints: the reliable authentication depends heavily on complex verification processes, expensive specialized equipment, and trained personnel; meanwhile, scalable production is hampered by by high costs, long cycle times, and environmental concerns. Although incremental progress has been made in specific technical areas, including encryption ^4,17^, encapsulation ^6,8,14,15^, and verification ^7,16,18^, the persistent, interrelated limitations collectively constitute a systemic barrier to broader implementation. Consequently, no DNA-based anti-counterfeiting technology to date has achieved a comprehensive and balanced breakthrough across the core metrics of security, convenience, cost-effectiveness, and environmental sustainability. InfinMark represents such a breakthrough that equips products with an invisible, irremovable, and encrypted taggant, providing reliable authentication regardless of the products’ physical form and even after its physical damage. Even under complex environmental stressors, the DNA fingerprint NPs remain stable and can be effectively extracted and identified throughout the product lifecycle, enabling long-term authentication and stricter supervision.

The InfinMark anti-counterfeiting system is designed to extend beyond encrypted coding and invisible tagging technologies, aiming to seamlessly integrate the real-time tracking of tangible anti-counterfeiting technologies, the the unclonable security of chemical unclonable functions (CUF) ^19^ and the immutable ledger of blockchain architecture, thereby establishing a multi-layered, bimodal encryption ecosystem geared toward future needs. The resulting multi-encrypted information is structurally non-reproducible and logical irreversible, rendering any attempt at decryption or replication via reverse engineering both technically and economically infeasible. Given its strengths in encryption intensity, concealment, material compatibility, and physical durability, InfinMark is poised for near-term deployment in fields with stringent anti-counterfeiting and traceability demands, such as high-value-added products, patent-intensive technology goods, and strictly regulated commodities. As DNA synthesis efficiency improves and sequencing costs decline, InfinMark is expected to accelerate the adoption of the “DNA ID for physical objects”. Its application is likely to expand from high-end applications into broader practical scenarios, including large-scale industrial manufacturing, farm-to-fork food traceability, pharmaceutical safety supervision, and personalized product customization. This heralds a next-generation IoT ecosystem,where a secure, unique identity is inherent to every object.

## Data availability

All data generated or analysed during this study are included in the main text or the supplementary table. The source data will be archived in a public repository upon publication. Prior to publication, the source data will be made available for peer review upon reasonable request.

## Code availability

Code for the encoding/decoding algorithm is available on GitHub at: https://github.com/qzds/Dynamic-Transcoding-Algorithm.

## Acknowledgements

This work was funded by National Key Research and Development Program of China (2024YFA0916503), National Natural Science Foundation of China (32122050 and 32370074) and Fujian Provincial Natural Science Foundation of China (2025J011005). We are grateful to Caiming Wu, Luming Yao and Wei Han from Core Facility of Biomedical Sciences, School of Life Sciences, Xiamen University for help in performing of electron microscopy.

## Author contributions

Z.Q.L. conceived and supervised the project, and together with T.H., designed the experiments, analyzed the results, and wrote the manuscript. B.Z.Z. developed the infoTrace kit, conducted all On-site Multi-Scenario Authentication experiments, and performed sequencing for all samples. T.H. and X.Z. developed the InfinMark nanoparticles (NPs) and completed all characterization and performance testing. F.K.H. and J.K.C. devised the codec algorithm and evaluated its performance, while Z.J.L. supervised the entire development process and contributed critical market-oriented insights. Z.M.Z. analyzed all sequencing data, while Y.F.G. calculated the half-life of InfinMark NPs under various temperature and humidity conditions. Y.F.G., L.L., and K.J.F. assisted in the development and testing of the InfinMark NPs encapsulation method. J.T. and S.N.L. were responsible for the synthesis of DNA sequences and the subsequent construction of plasmid vectors. A.N.L., X.Z.K., H.W., Q.Y.Z., and Z.Y.Y. provided technical guidance and assisted in manuscript preparation.

