## Supplementary information for "End-to-end bimodal anti-counterfeiting by informational DNA nanoparticles"

### Methods

#### 1. Cryptographic encoding and decoding of anti-counterfeiting information

**Encoding.** To address the short-text characteristics of anti-counterfeiting traceability information, we designed a DNA encoding algorithm based on character compression and triplet base mapping. First, the input string (containing 10 digits and 26 English letters) is scanned for repeated sequences. When consecutive identical characters exceed a length of 5, their position and length information are extracted as metadata, and the original string is compressed. The position and length information in the metadata are then converted into base-36 strings, concatenated in a "position + length" format, and terminated with a 'z' character. Both the compressed data and metadata are encoded using randomly generated triplet base mapping tables, where a bijective relationship is established between the 36 characters and 36 triplet nucleotide combinations that avoid consecutive identical bases. To optimize biochemical sequence properties, the algorithm generates candidate sequences through multiple rounds of random search and employs a comprehensive scoring system based on GC content (40%–60%) and total homopolymer length, ultimately selecting the sequence with the highest composite score as the encoding result.

**Decoding.** A complete decoding workflow is designed to recover the original information from encoded DNA sequences. First, the system validates that the input sequence contains only A, T, C, and G bases to ensure data integrity. Then, it loads the corresponding JSON mapping file, which records the metadata flags and mapping relationships used during encoding. If the sequence contains no metadata (has_meta = 0), it directly divides the sequence into triplets and decodes each triplet to original characters using the reverse mapping table. If metadata is present (has_meta = 1), the algorithm first decodes the metadata section: it processes triplets sequentially into base36 characters until encountering the terminator 'z', then converts pairs of base36 strings into position indices and repetition lengths to generate a list of duplicate units. The remaining sequence portion is similarly decoded via triplet mapping into a compressed string, which is finally expanded according to the duplicate unit list to restore the complete original text by reconstructing the repetition patterns.

**Evaluation.** To validate the effectiveness of our encoding algorithm, we collected real-world data including tea varieties, production regions, and harvest times from public sources. After converting this information to Pinyin and extracting initial letters for concatenation, we constructed a test dataset containing 19,951 entries. Using the standard workflow from the Chamaeleo toolkit, we compared our algorithm with mainstream DNA storage encoding methods including Church, Goldman, Grass, Blawat, and DNA Fountain.

**2. Synthesis, construction and extraction of anti-counterfeiting DNA fingerprints for encapsulation.** The encrypted DNA sequence carrying anti-counterfeiting information of Xiamen University motto "ziqiangbuxizhiyuzhishan" was constructed into the SUP149 plasmid. Single-stranded oligonucleotides (Oligos) with overlapping regions (see Supplementary Table 3) were used for assembly of the anti-counterfeiting information-containing DNA fragments via the PCA method [20]. The resulting fragments were then cloned into pRS415 using the Cloning Kit (SC612, Genesand Biotech Co., Ltd.), followed by transformation into *E. coli* DH5α strain for plasmid cloning. The plasmid was extracted and purified using the Alkaline Lysis Method [21]. The three lowest-scoring tea anti-counterfeiting DNA fingerprints were produced by GENEWIZ Biotech Co., Ltd., with supporting mass spectrometry analysis reports.

To develop lowcost anti-counterfeiting DNA fingerprint nanoparticles, we propose a bio-manufacturing strategy using engineered cellular systems: such as *E. coli* (for short sequences, bp-kb level) and *S. cerevisiae* (for long sequences, 30 kb-Mb level). This approach allows industrial-scale synthesis using only a 5-liter fermenter.

**3. Preparation of anti-counterfeiting DNA nanoparticles.** DNA fingerprints were prepared as a 100 ng/µL aqueous solution. The 10% (w/v) gelatin and 2% (w/v) chitosan stock solutions were prepared by dissolving the powder in a 40°C water bath (gelatin) or on a shaker of 37 °C (chitosan), respectively. Gelatin working solution consisted of 1.5 mL 10% stock, 1.5 mL glacial acetic acid, and 10 mL MQ water, and chitosan working solution contained 7.5 mL 2% stock, 1.5 mL glacial acetic acid, and 4 mL MQ water. For DNA encapsulation, 18 µg of plasmid DNA was added to 6 mL of working solution and incubated for 30 min on shaker. After the addition of 3 mL 25 wt% crosslinker (glutaraldehyde or vanillin), the reaction mixture was agitated continuously for 4 h to ensure complete crosslinking. The resulting DNA-loaded microspheres were collected by centrifugation after pH adjustment and washed twice with MQ water. Silica encapsulation was performed by reacting the microspheres with dilute sodium silicate (50-fold dilution of saturated solution, adjusted to pH 10 with acetic acid) for 24 h.

**4. Nanoparticle characterization.** The particle size distribution, dispersity and Zeta potential of the encapsulated DNA were measured using a Malvern Panalytical Zetasizer Ultra (Malvern Instruments Ltd., Malvern, UK) at 25.0 °C. Encapsulated DNA fingerprint NPs were diluted to an appropriate concentration, gently vortexed, and equilibrated for 3 min prior to analysis. For zeta potential measurements, the suspensions were analyzed following application of a 20 V/cm electric field and converted into the zeta potential according to the Helmholtz-Smoluchowski equation. Data are presented as mean ± SD from three independent experiments. For SEM imaging, the nanoparticles were diluted with MQ water to an appropriate concentration. A 20 µL droplet of the suspension was deposited on a silicon wafer and air-dried. The sample was then sputter-coated with a 5–10 nm platinum film and examined by SEM at an accelerating voltage of 5 kV. High-resolution micrographs were acquired at magnifications between 10,000× and 100,000× by fine-tuning the working distance, objective aperture, and astigmatism correction. For nano-flow cytometric analysis, DNA was stained with SYBR Green I at dye-to-sample ratios of 1:10,000 and 1:2,000 for the gelatin and chitosan nanoparticle formulations, respectively. The staining was performed for 30 min at room temperature in the dark; excess dye was then washed away prior to silica encapsulation. To avoid interference from aggregates, all prepared nanoparticle samples were filtered through a 0.45 µm membrane before analysis. Using the 488 nm laser for excitation, the instrument was configured with a side scatter attenuation coefficient (ssDcay) of 10% and laser power of 10 mW for gelatin NPs, and 0.2% ssDcay and 20 mW for chitosan NPs.

**5. Stability under harsh conditions.** Aqueous suspensions of naked DNA or DNA NPs were subjected to three distinct mutagenic/degradative challenges. For UV exposure, samples in uncapped microcentrifuge tubes were placed 10 cm from the UV source (254 nm, 100 mW/cm$^2$) and irradiated for 1, 2, 3 and 4 h. For freeze-thaw stress, samples underwent 5, 10, 20, or 30 cycles, with each cycle consisting of freezing at -80°C for 30 min and thawing at 37°C for 5 min. For oxidative stress, NP suspensions were treated with $H_2O_2$ at final concentrations of 5, 10, 15, 20,

25, 30, and 35 mmol/L for 1 h at 25°C, followed by heat treatment to inactivate residual $H_2O_2$. The DNA retention rate was analyzed by qPCR and normalized to that of the untreated controls.

**6. Thermal aging test.** Lyophilized samples of naked DNA and DNA NPs were prepared and placed in sealed containers with reservoirs of saturated salt solution ($MgCl_2$, NaBr, and NaCl), to keep the relative humidity level at 25%, 50% and 75%, respectively. The sealed containers were placed in laboratory ovens maintained at 55°C, 65°C, or 75°C. All samples were aged concurrently across the full matrix of temperature and humidity conditions. Replicate samples were collected at predetermined intervals (0, 1, 2, 3 and 4 weeks). Following the aging period, all samples were rehydrated in nuclease-free water, and the remaining DNA was quantified by qPCR. Half-life values were calculated by applying the Arrhenius equation, assuming first-order decay kinetics, as described previously[11].

**7. Tagging and extraction of anti-counterfeiting DNA nanoparticles in multiple materials.** (1) Incorporation into ink. To prepare the DNA-tagged stamp ink, anti-counterfeiting DNA NPs (0.002 g) were dispersed in 1 mL of commercial ink and homogenized on a rotary platform for 2 h. A 10 μL aliquot was sampled for quality control before the remaining formulation was applied to a photosensitive stamp. After stamping, the imprint was air-dried for 15 min. DNA fingerprint NPs were then sampled from the dried imprint by gentle swabbing with a cotton bud, followed by alkaline lysis in 100 μL of 0.2 M NaOH, releasing the embedded DNA for subsequent analysis. (2) Incorporation into gunpowder. Gunpowder was harvested from commercially sourced firecrackers (GongXiFaCai 0.7 Series) by dissecting the paper packaging. The collected gunpowder was then blended with encapsulated DNA NPs at a ratio of 0.002 g per 1 g of powder. DNA recovery was evaluated both before and after combustion. For pre-combustion analysis, 50 mg of the uncombusted mixture was added to 100 μL of 0.2 M NaOH. For post-combustion analysis, one-tenth of the ash obtained from combusting 0.5 g of the mixture was treated with 100 μL of 0.2 M NaOH to release the DNA. (3) Incorporation into PCL plastic. A 0.002 g quantity of DNA NPs was blended with 1 g of commercial polycaprolactone (PCL) particles. The mixture was heated on a 100°C hot plate to melt the PCL. The resulting viscous liquid was stirred thoroughly in its molten state to ensure complete homogenization before solidification at room temperature. Approximately 0.05 g of the solidified composite was sectioned and and incubated in 100 μL of 0.2 M NaOH to facilitate subsequent DNA release.

**8. Rapid on-site product authentication**

**Sample pretreatment.** An appropriate amount of the test sample was collected using a swab or tweezers and transferred into Tube L1. After thorough mixing, the tube was subjected to five cycles of thermal treatment (3 min at 95°C followed by 2 min at 4°C). Subsequently, 100 μL of Buffer L2 was added to the Tube L1, mixed thoroughly, and the resulting lysate was reserved for downstream use.

**Specific amplification of anti-counterfeiting DNA fingerprint.** The lyophilized powder in Tube S1 was first fully resuspended with 40 μL of Buffer S2 (from Tube S2). Then, 5 μL of the pretreated alkaline lysate from the previous step and 5 μL of Buffer S3 were added to Tube S1, and the mixture was thoroughly vortexed. The reaction was incubated at 30°C for 15 min to allow amplification. After incubation, 1 μL of the amplification product was transferred into Tube S4 for dilution and reserved for subsequent analysis.

**Rapid colorimetric identification of authenticity.** 25 μL of Buffer K1 was transferred from Tube K1 into Tube G and mixed thoroughly. Subsequently, the diluted amplification product from Tube S4 (previous step) was added and mixied uniformly. The color change of the reaction mixture was observed at room temperature to determine the presence of the anti-counterfeiting DNA fingerprint in the sample. For validation, a parallel negative control reaction using pure water is recommended. All the above components and their functions are detailed in Table S7.

**9. Amplification, sequencing and recovery of anti-counterfeiting DNA fingerprints.** Following the addition of 0.2 M NaOH, all samples were subjected to thermal cycling (3 cycles of 98°C for 3 min and 4°C for 2 min), followed by neutralization with 0.2 M HCl. Neutralized lysate was then used as the template for PCR (TransGen AP111-01), or InfoTrace kits. For sequencing library construction, a two-step PCR amplification method was employed. First, the alkaline lysis products served as templates for amplification with primers seq1F and seq1R, which contained unique barcode sequences. The resulting PCR products were gel-purified, and equal amounts were used as templates for the second PCR amplification with primers BZO426 and BZO427. All primer sequences used for amplification are detailed in Table S8. The final PCR products were purified by gel extraction and submitted to Biomarker Technologies for sequencing on an Illumina Nova6000 PE150 platform. To analyze the sequencing data, the original read order was maintained. In each reconstruction run, 250 reads were randomly selected as starting points. Beginning from each start point, every other read (i.e., the 1st, 3rd, 5th, 7th, … read in sequence order) was included until the 999th read was reached. The nucleotide frequency was then calculated for each position across all covering reads, and the most frequent base was assigned as the consensus to complete sequence assembly. The assembled sequence was aligned to the reference to compute its recovery rate. The final evaluation metric was the average recovery rate across all 250 starting points.

**10. Cost-Performance-Index (CPI) calculation**

To quantify the comprehensive efficacy of InfinMark as a security product, this study introduces a cost-performance index (V), defined as a function of effective security bits, protection duration, and unit cost:

$V = (\log_2 S \cdot w \cdot T) / C.$

This model integrates four dimensions—cryptographic space (S), engineering-experience weighting (w), durability (T), and cost (C)—to enable an objective and repeatable quantitative comparison of the overall performance of different anti-counterfeiting technologies.

In the formula:

Effective security bits are derived from $\log_2$ (cryptographic space S) and the encryption-level weight w. The value of S is obtained from published literature and practical calculations. As noted in the literature [22], taking the logarithm converts cryptographic spaces of different bases into a uniform "bit-length" metric. This allows the strengths of various encryption techniques to be compared on a common scale, preventing the disproportionately large absolute value of DNA cryptographic space from overshadowing the contributions of lifespan (T) and cost (C). The weight w serves as an empirical "quality factor" that differentiates the practical difficulty of breaking the security under varying technical barriers: publicly visible to the naked eye (L1, w = 1), requiring tools/specialized devices (L2, w = 1.5), and requiring laboratory-grade analysis (L3, w = 2). This design ensures that the effective security bits reflect both algorithmic strength and implementation barriers, addressing the limitation of "comparing only bit length while ignoring practical robustness". The protection duration T refers to the average number of years the anti-counterfeit feature remains effective on the product, which can be approximated as the label's lifespan. Its value is sourced from literature and experimental data. The unit cost C is based on actual production accounting and market research data.

**11. Statistical information**

All statistical analysis were performed using unpaired two-tailed Student's t-test in GraphPad Prism 8.0 software. Detailed statistical information, including exact sample sizes (n), is provided in the respective figure legends. Significant differences of all results are indicated as *$p < 0.05$, ** $p < 0.01$, and *** $p < 0.001$; ns, no significance. Investigators were not blinded to group allocation during experiments.

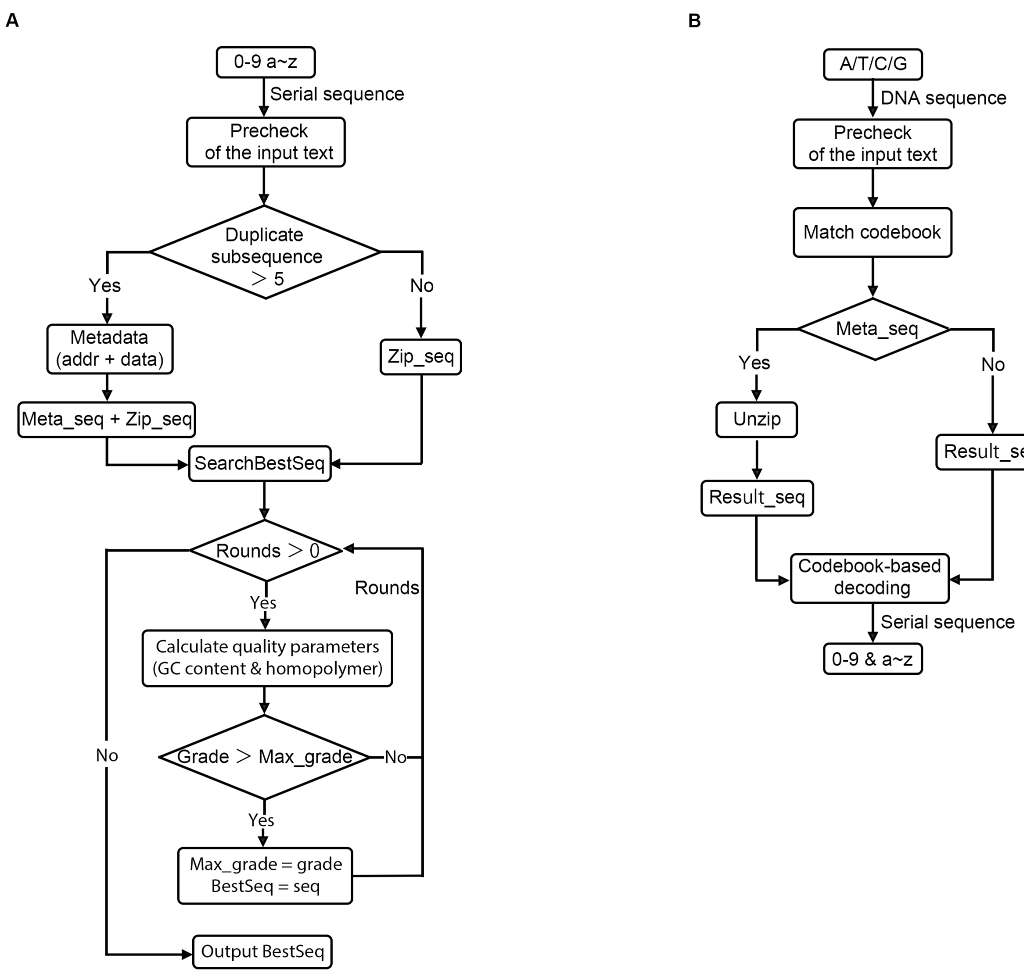


**Figure S1. Workflow for cryptographic encoding and decoding of anti-counterfeiting information. A.** Schematic of the procedure for encrypting and encoding anti-counterfeiting information into a DNA sequence. The process begins by verifying input character legitimacy and scanning for long repetitive sequences (>5 repeats). Metadata compression is then applied, where positions and counts of repetitive subsequences are extracted to generate metadata, and the original string is deduplicated to increase encoding density. The validated and compressed data are then dynamically encoded. This involves converting characters to DNA bases via randomly generated mapping tables, followed by iterative optimization. Candidate sequences undergo multi-round screening against biochemical constraints (e.g., GC content, homopolymer length). The sequence with the highest composite score is selected as the final output (BestSeq). **B.** Decoding workflow. The input DNA sequence is first validated to contain only A, T, C, and G. The corresponding codebook is then identified. If a metadata flag is present, a decompression step is initiated: repeat locations and counts are extracted from the metadata and used to reconstruct the original sequence by restoring the repeated segments. Finally, the DNA sequence is mapped back to characters using the codebook, outputting the decrypted anti-counterfeiting and traceability information in plaintext.

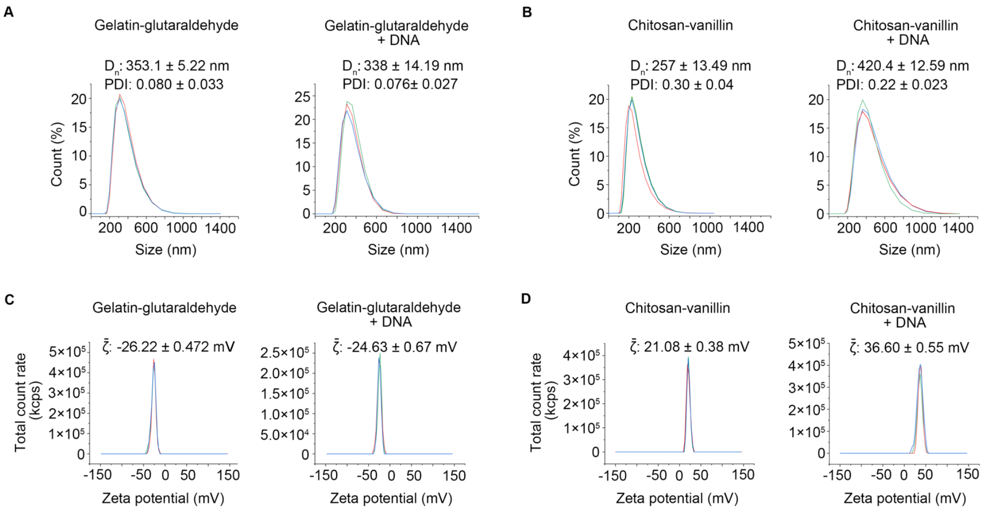


**Figure S2. Size distribution, dispersity and zeta potential of anti-counterfeiting DNA NPs. A-D.** Gelatin-glutaraldehyde and chitosan-vanillin nanoparticles, with and without encapsulated DNA fingerprints, were diluted 100-fold for particle size (**A, B**) and zeta potential (**C, D**) measurements. Data are expressed as mean ± SD. Dn, PDI, and $\zeta$ represent the number mean diameter, polydispersity index, and mean zeta potential, respectively. Mean values from n = 3 independent experiments.

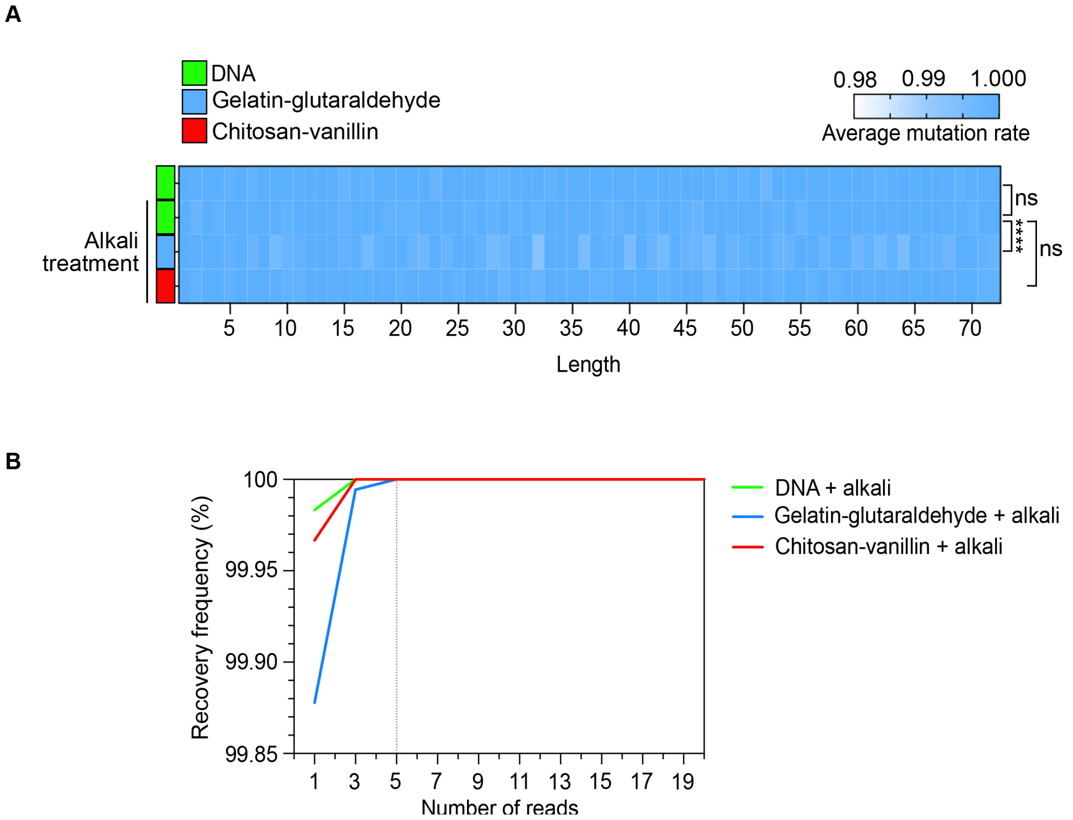


**Figure S3. Effect of encapsulation and alkaline decapsulation on DNA fidelity. A.** Naked DNA, gelatin-glutaraldehyde- and chitosan-vanillin-encapsulated DNA fingerprints were subjected to alkaline treatment followed by Illumina sequencing. The heatmap shows the per-base mutation rate across the 72-bp encoding region, derived from the first 1,000 sequencing reads. **B.** Recovery rate as a function of sequencing depth. The relationship was assessed by analyzing random read subsets. For each of 250 random start points, every other read was selected (e.g., reads 1, 3, 5, …) sequentially until the 999th read was included. The curve shows the mean recovery rate across all 250 subsets at each sequencing depth.

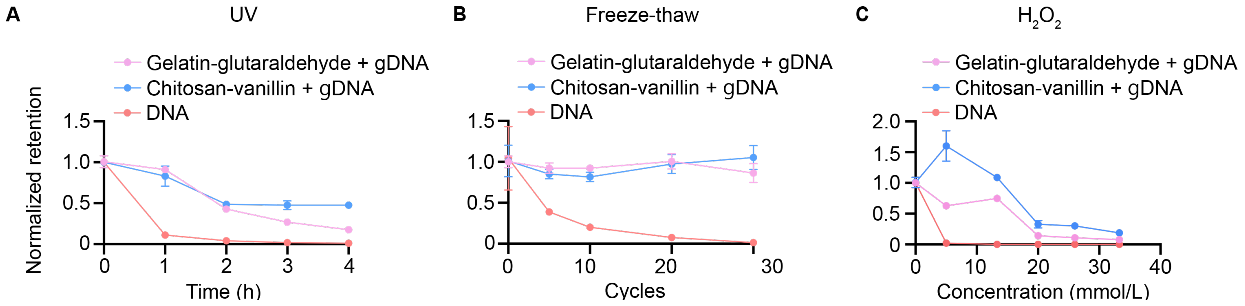


**Figure S4. Encapsulation enhances the retention rate of anti-counterfeiting DNA fingerprints under harsh conditions.** DNA retention rates were measured using qPCR for naked, gelatin-glutaraldehyde-, and chitosan-vanillin-encapsulated DNA fingerprints under the following stresses: (A) varying durations of UV irradiation at 10 cm; (B) multiple freeze-thaw cycles between 37°C and -80°C; (C) increasing concentrations of $H_2O_2$. All bars in the figure represent mean ± S.E.M., n = 3 independent samples.

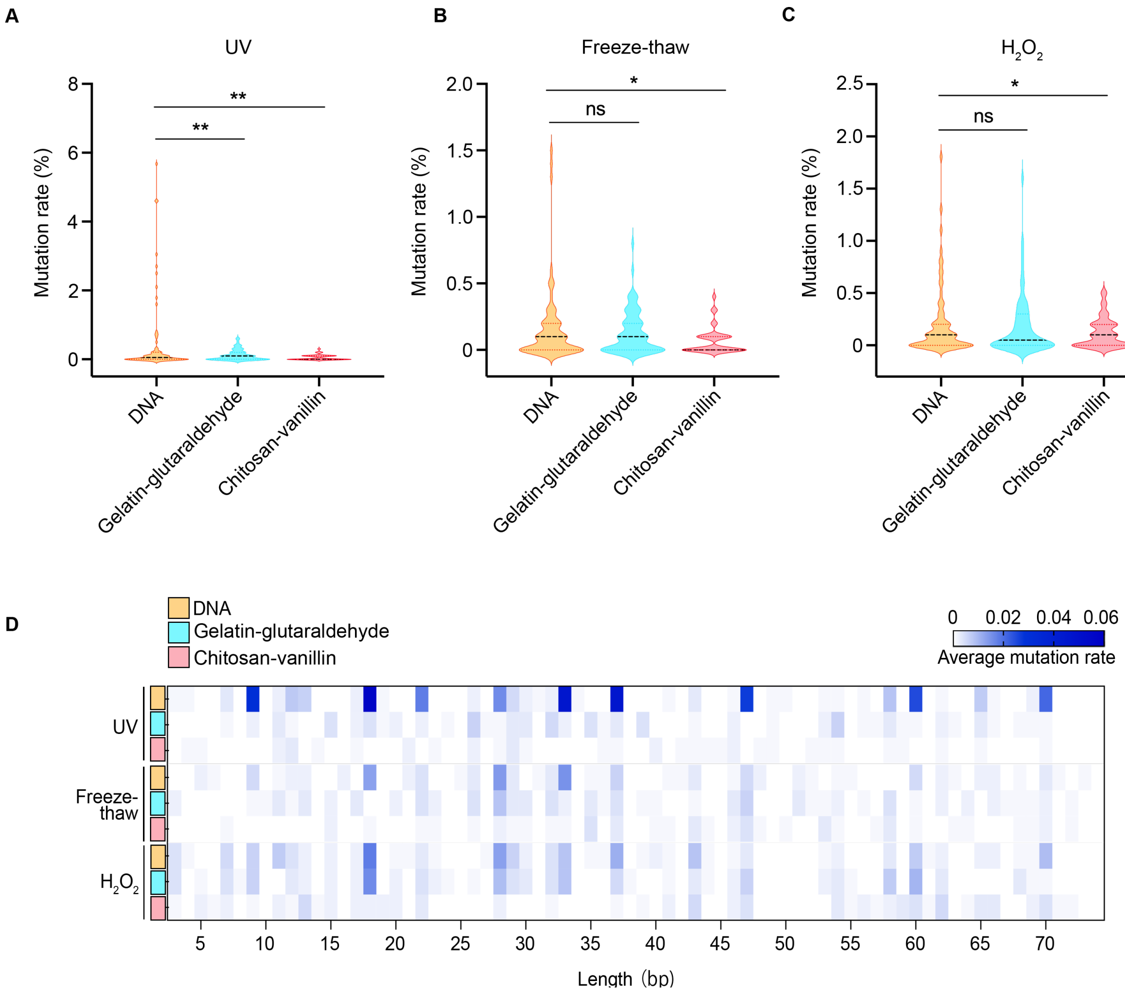


**Figure S5. Encapsulation reduces the insertions-deletions-substitutions (IDS) mutation rate of anti-counterfeiting DNA fingerprints under harsh conditions. A-C.** IDS mutation rates under stress. IDS mutation rates were measured by Illumina sequencing and analyzed for naked, gelatin-glutaraldehyde-, and chitosan-vanillin-encapsulated DNA fingerprints under the following stresses: (**A**) exposed to ultraviolet light at a distance of 10 cm for 20 minutes; (**B**) 5 freeze-thaw cycles between 37°C and -80°C; (**C**) exposed to 1.6 mM $H_2O_2$ for 1 hour. (**D**) Per-base mutation profile. The heatmap displays the average mutation rate at each position across the 72-bp encoding region, calculated from the first 1,000 reads.

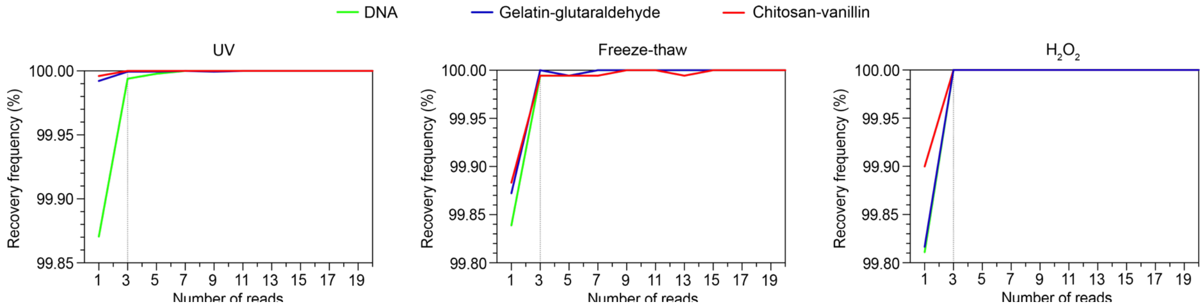


**Figure S6. Sequencing depth required for accurate recovery of DNA fingerprints under varying harsh conditions.** Naked, gelatin-glutaraldehyde-, and chitosan-vanillin-encapsulated DNA fingerprints were subjected to the following stresses: (**A**) ultraviolet exposure for 20 min at a 10 cm distance; (**B**) 5 freeze-thaw cycles between 37°C and -80°C; and (**C**) treatment with 1.6 mM $H_2O_2$ for 1 hour. For each condition, the minimum sequencing depth required to achieve accurate sequence recovery was determined from the Illumina sequencing data.

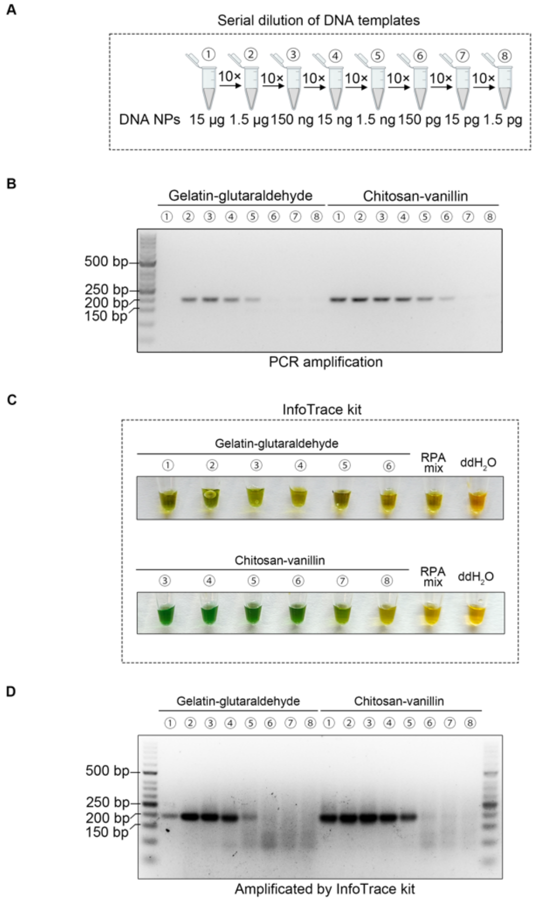


**Figure S7. Performance of the InfoTrace kit in identifying anti-counterfeiting DNA NPs**. **A.** Serial dilution of nanoparticles. Gelatin-glutaraldehyde- and chitosan-vanillin-encapsulated DNA fingerprint NPs were subjected to ten-fold serial dilution (nanoparticle weight shown; DNA content is substantially lower). **B-C.** Detection sensitivity. At each dilution, DNA was extracted and analyzed by conventional PCR (**B**) and the InfoTrace rapid detection kit (**C**). **D.** Gel verification of InfoTrace kit. Amplification products from the InfoTrace kit were confirmed by agarose gel electrophoresis. The assay revealed a detection limit of 1.5 ng for gelatin-glutaraldehyde nanoparticles and 15 pg for chitosan-vanillin nanoparticles.

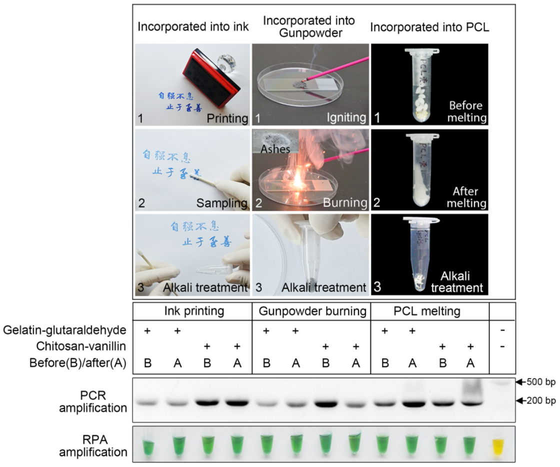


**Figure S8. Authentication of DNA fingerprint in practical applications.** DNA fingerprint nanoparticles were incorporated into stamp ink, gunpowder, and PCL. Following exposure to their respective stresses (stamping, combustion, or melting at 100°C), DNA was successfully detected by both PCR and the InfoTrace kit. "B" and "A" denote samples before and after stress, demonstrating robust authentication across all scenarios.

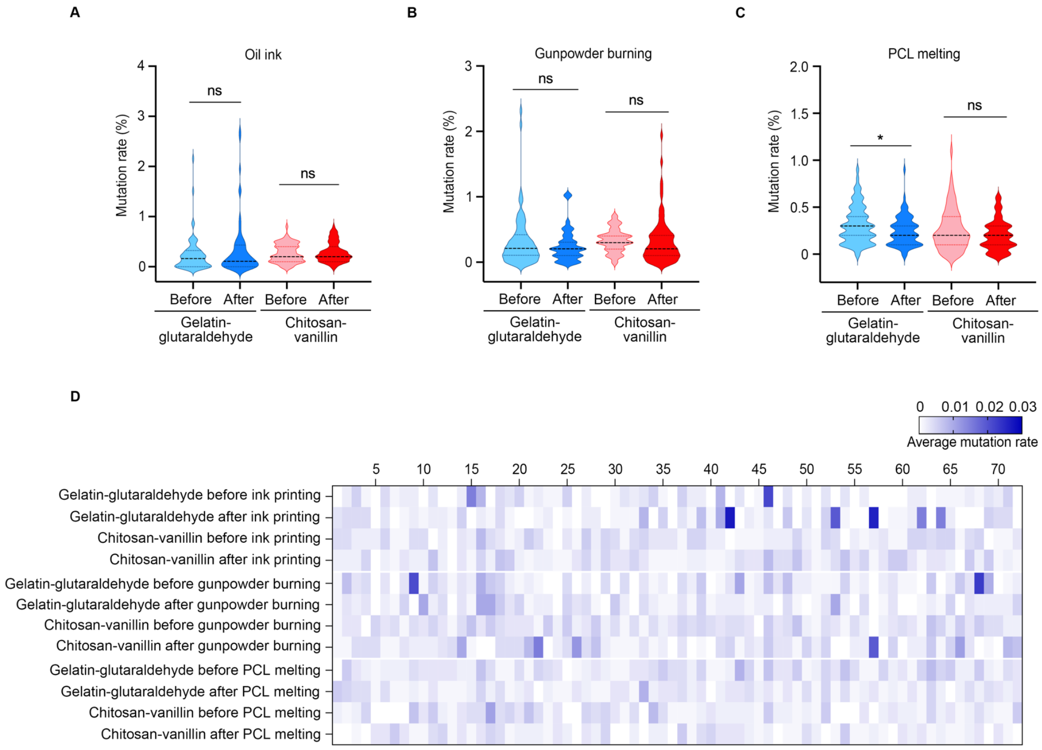


**Figure S9. IDS mutation rate of DNA fingerprint NPs in practical applications. A-C.** Mutation rates after physical stress. DNA fingerprint nanoparticles incorporated into oil ink, gunpowder, or PCL were sampled before and after (**A**) stamping, (**B**) combustion, or (**C**) melting at 100°C. Insertion-deletion-substitution (IDS) mutation rates were analyzed using Illumina sequencing data. **D.** Per-base mutation profiles. The heatmap shows the average mutation rate at each position across the 72-bp encoding region for each experimental condition.
